# Comparative Genomics of the Lipid Droplet-Associated Protein Seipin Across Eukaryotic Diversity Illuminates An Ancient Origin and Conserved Structural Diversity

**DOI:** 10.1101/2025.10.20.683490

**Authors:** Emily M. Kingdon, Muskaan Kainth, Hareem Tariq, Fiona Karaj, Elisabeth Richardson

**Affiliations:** Mount Royal University, Calgary, Alberta, Canada; University of Alberta, Edmonton, Alberta, Canada

**Keywords:** Lipid Droplets, Bioinformatics, Comparative Genomics, Evolutionary Cell Biology, Protistology

## Abstract

Lipid droplets (LDs) are ubiquitous across living organisms. Characterised in eukaryotes by their lipid monolayer and essential for lipid storage and metabolism in animals, plants, and yeast, little is known about the evolutionary diversity of extant LDs across the eukaryotic tree of life. LDs facilitate an impressive variety of cellular functions outside of just lipid storage; in Metazoa, these include stress responses, cellular signaling, and membrane remodeling. Likewise, the distribution of these functions across eukaryotic diversity is unknown.

We have examined the evolutionary trajectory of seipin, a protein associated with LD biogenesis, across eukaryotic diversity. We have identified a pan-eukaryotic distribution of seipin, with an evolutionary pattern that indicates presence in the Last Eukaryotic Common Ancestor. Ancient conservation of multiple variants of seipin suggests that seipin may multiple conserved functional roles within the cell. Finally, we identify a lack of sequence homology between BSCL2/seipin in *Homo sapiens* and the *sei-1* gene previously identified as a functional homologue of BSCL2 in *Saccharomyces cerevisiae*. Though our results suggest that *sei-1* may be a highly divergent homologue of BSCL2 / seipin rather than an unrelated protein, the divergence suggests that researchers may need to be cautious applying results obtained in *Saccharomyces spp.*.

**Visual abstract:** 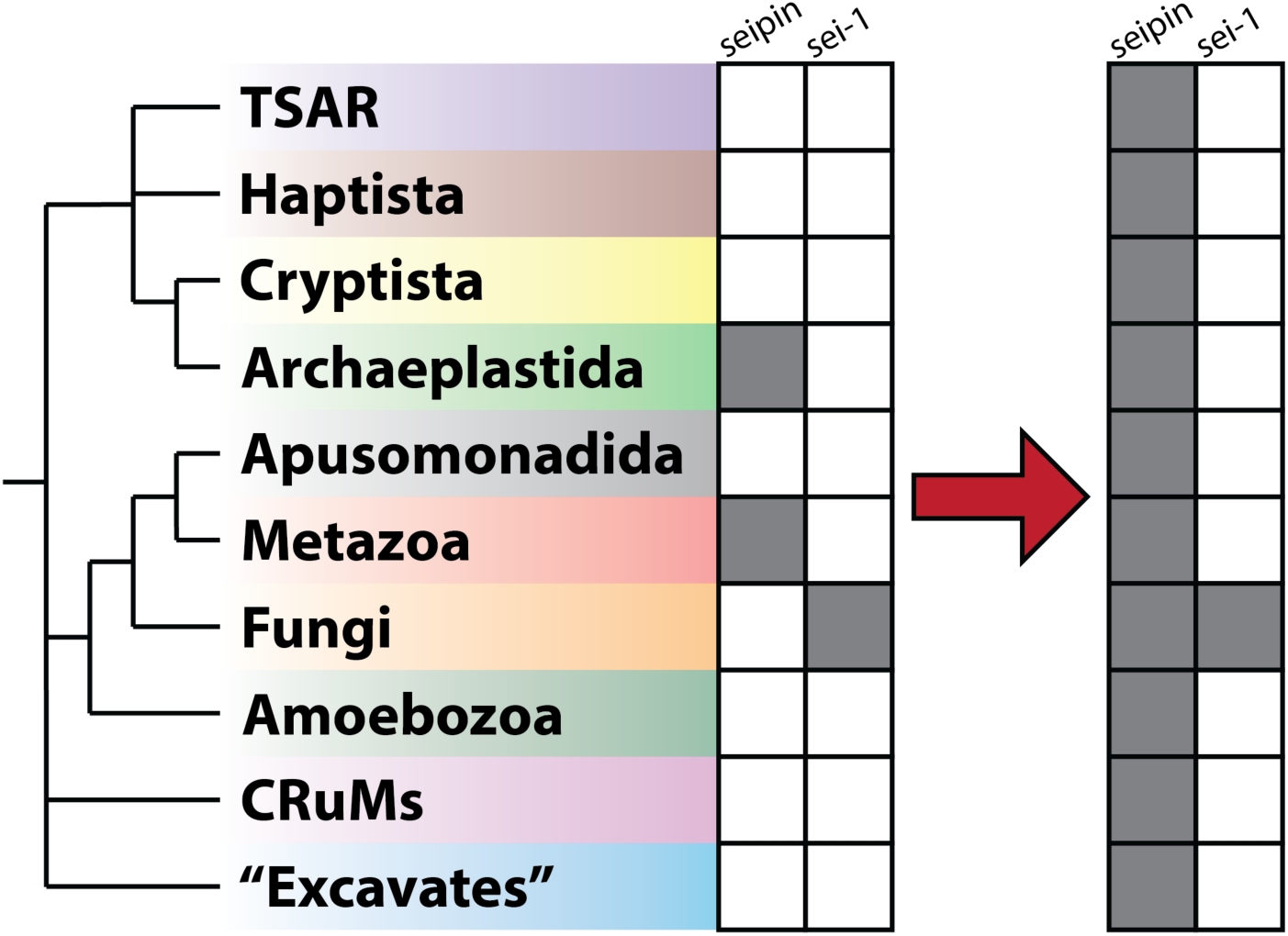

## Introduction

Long described as “enigmatic”, lipid droplets (LDs) are a ubiquitous organelle in prokaryotes and eukaryotes alike^1^. They have long been studied in humans and related model organisms due to their role in human health and disease; mutations in LD-associated proteins (LDAPs) have been implicated in heritable genetic conditions such as lipodystrophy and various neurodegenerative syndromes^2^. Our understanding of lipid droplet biology in metazoan model systems is advanced, and medical researchers have identified roles for LDs and LD-associated proteins in including stress responses^3^, cellular signalling^4^, membrane remodeling^5^, and mitigating the effects of reactive oxygen species (ROS)^6^. In these systems, LDs have been found to form *de novo* through the slow buildup of neutral lipids in the endoplasmic reticulum and retain phospholipids from the outer leaflet of the ER membrane even after division^7^, although biogenesis of LDs via binary fission has been identified in *Saccharomyces cerevisiae*^8^. LDs can associate with most other cellular organelles through membrane contact sites^9^, which are dynamic and adapt to periods of LD expansion and shrinkage. The functionality of LDs is proposed to be dictated by their proteome^10^, which in mice has been found to contain lipid metabolic enzymes, regulatory scaffold proteins, proteins involved in LD clustering and fusion, as well as other proteins with unknown functions^11^. LD biogenesis and degradation, as well as their interactions with other organelles, appears to be tightly coupled to cellular metabolism in multicellular organisms. This relationship, in turn, relates to the maintenance of lipotoxicity^9,12^. The dynamic nature of LDs and their interactions with other cellular components highlight their importance in maintaining cellular lipid homeostasis.

In parallel, an equally compelling story of LD function has emerged in plants and algae, mostly due to research into biofuels and carbon storage^1^. Studies in plant and algal models such as *Arabidopsis thaliana* and *Chlamydomonas reinhardtii* have also revealed roles for LDs in pathogen detection and immune response^13^, development of seeds^14^, and response to environmental contamination^15^. During periods of stress, there is an observed increase in the presence of LDs in the leaves and microalgae of plants and algae respectively^16,17^. The model green alga, *C. reinhardtii* displays an experimentally well-documented upregulation of cytoplasmic LDs upon transition to nitrogen-depleted growth medium, which pushes the cells into a dormant state waiting for more favorable growth conditions^1,17^. Algal LDs are also being explored as a source of biofuels^18^.

Comparing the metazoan and archaeplastid research literature, we find that LDs share a common ultrastructure, with a single phospholipid bilayer enveloping a hydrophobic neutral lipid core (Figure 1A). However, their diversity of composition, morphology, function, and sub-cellular localization is as expansive as their pan-eukaryotic distribution^1,19^. Initially, LD biogenesis was considered a purely biophysical process akin to the separation of oil and water. However, genomic, metabolic and proteomic studies have also identified a variety of genes that are associated specifically with LDs and are essential for LD localisation, structure, and function^10^. Notably, families of genes associated with biogenesis and maintenance of LDs, such as the perilipins in Metazoa and the oleosins in Archaeplastida, appear to be restricted to organisms within and closely related to the lineages in which they were originally identified^1^. This is unusual, as highly conserved membrane-bound organelles are usually accompanied by a conserved proteome and this raises questions on the origins and evolutionary trajectory of LDs as an organelle^20^. Either the biophysical necessity of a dynamic organelle for lipid storage has resulted in the convergent evolution of multiple organelles with LD characteristics, or an ancient LD-like organelle evolved in the Last Eukaryotic Common Ancestor (LECA) and has been retained for its ancestral function of lipid storage throughout eukaryotes, with additional functionality evolving over time. These lineage-specific functions may be associated with the previously identified lineage-specific protein families like oleosins and perilipins^21^.

**Figure 1.**
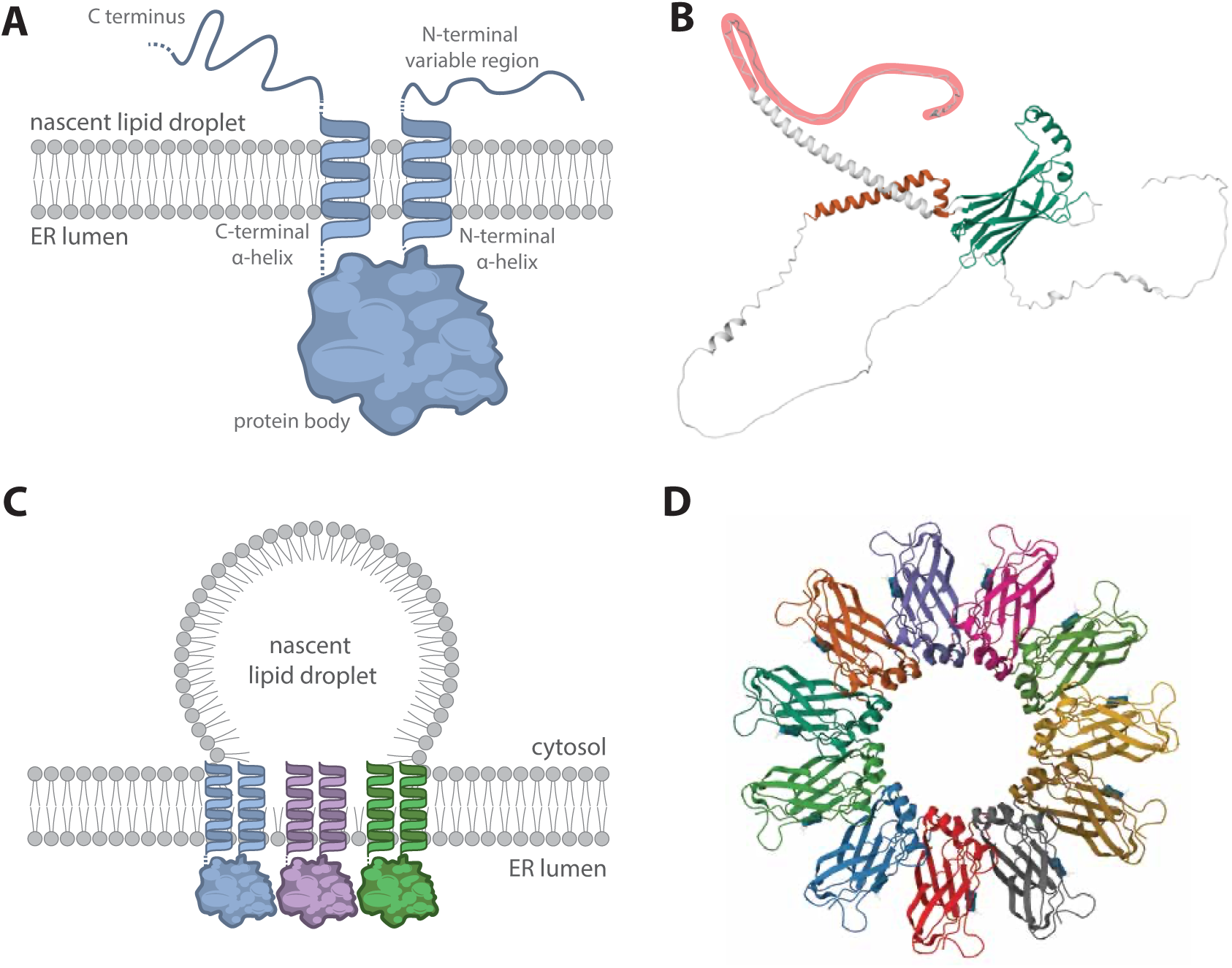
Involvement and ultrastructure of seipin in the formation of lipid droplets. A) Seipin has two transmembrane domains (TMD) which cross the lipid bilayer between the ER lumen and a forming nascent lipid droplet. This consists of the C-terminal region in the nascent lipid droplet, followed by the C terminal alpha helix TMD. Within the ER lumen is the protein body, which then exits using the N-terminal alpha helix TMD, followed by the variable N-terminal region back out into the nascent lipid droplet. B) The protein structure of a single seipin subunit, derived from an AlphaFold predicted structure of NP_001372957.1, a *Homo sapiens* isoform of seipin / BSCL2 which includes the extended N terminus (highlighted in red). C) Schematic of a lipid droplet forming from the ER membrane and pinching off from the lipid droplet, showing the position and orientation of seipin (not to scale). Adapted from Gao et al (2019)^7^. D) Seipin copies cluster together to form the transmembrane basket structure which serves to pinch off lipid droplets during formation. This image is derived from the PDB structure 6DS5, submitted by Yan et al (2018)^22^.

Seipin is a transmembrane protein essential for LD biogenesis from the ER, forming a polymeric basket-like structure around the base of the budding lipid droplet and contributing to its fission from the ER membrane (Figure 1A)^22,23^. Known as BSCL2 in mammals, it is also associated with metabolic and neurodegenerative disorders^2^. One of the identified linked diseases is congenital lipodystrophy, a rare autosomal recessive disease characterized by a near-absence of adipose tissue from birth or early infancy and severe insulin resistance^24^. Mice without seipin were shown to be growth delayed, but eventually achieved normal weight^25^. These mice showed significantly reduced adipose tissue mass, including about 60% reduction in brown adipose tissue, glucose intolerance, hyperinsulinemia, and hepatic steatosis, with elevated expression of select lipogenic genes^25^. In yeast, a protein known as Fld1p or sei-1 has been identified as a functional homologue of seipin^26^. This protein is also associated with LD biogenesis and structure, and has the TM-loop-TM structure conserved between all characterised homologues of seipin^23^. Since this protein has been found in organisms across the eukaryotic tree^1^, it is an ideal candidate for identifying initial patterns of LD evolution and diversification across the eukaryotic tree of life.

In this study, we use a comparative genomics approach to examine the evolution of seipin across eukaryotic diversity, as it is one of the few LDAPs known to have a pan-eukaryotic distribution. We have confirmed an ancient origin of seipin, likely within the Last Eukaryotic Common Ancestor (LECA), and that LECA likely contained multiple structural variants of seipin. This suggests that the additional roles of LDs outside of lipid storage identified in restricted lineages may have origins deeper in the evolutionary history of eukaryotes. Furthermore, we also observed that the functional homologue of seipin in *Saccharomyces cerevisiae*, the most used model unicellular eukaryote, does not appear to retain sequence homology with the other canonical homologues of seipin, though other fungi within the Ascomycota do have canonical seipin homologues. Phylogenetic analysis suggests that while sei-1 does share some relationship with pan-eukaryotic seipin, it is unusually divergent from the same protein in other Opisthokonta like humans and mice. This result suggests that sei-1 has undergone rapid diversification specifically within the Saccharomycetes, and any results obtained from *S. cerevisiae* models must be cautiously applied outside of this specific fungal class.

## Results

### Distribution of seipin across eukaryotic diversity

We assessed seipin distribution across eukaryotic diversity using queries from model organisms in which seipin has been characterised (Table 1), homology searching into selected genomes from organisms across eukaryotic diversity (Table S1). We combined a manual (Table S2) and automated (Table S3) approach to holistically detect even distant homologues while also allowing high-throughput homologue detection. The combined distributions are summarised in Table S4 and Figure 2. We detected a patchy but consistent distribution of seipin across eukaryotic diversity, with confirmed identification of at least one seipin homologue in 43 of the 78 analysed genomes (55.1%) and putative homologues in 53 of the 78 analysed genomes (70.0%) (Figure 2). Seipin homologues were also identified in 12/12 (100%) of analysed eukaryotic supergroups (Figure 2). This substantially expands the previously known distribution of seipin across eukaryotes; the gene had been described in detail in *Arabidopsis thaliana*, *Saccharomyces cerevisiae*, and various Metazoa including *Homo sapiens*, *Mus musculus*, *Drosophila melanogaster* and *Caenorhabditis elegans*. Though previous reviews of seipin distribution had speculated that this gene had a pan-eukaryotic distribution^1^, this is confirmed by our analyses.

**Figure 2.**
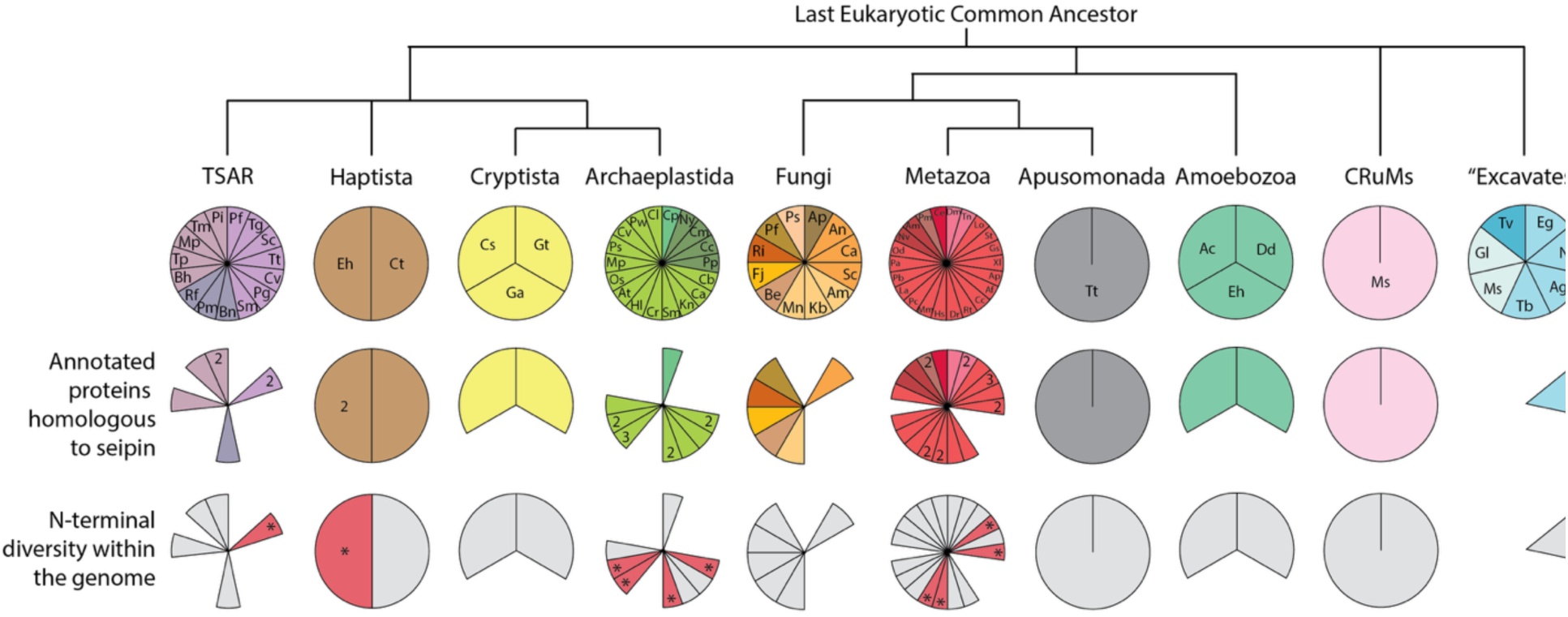
Coulson plots depiction of seipin homology and n-terminal variance among eukaryotic organisms, summarizing the information in Tables S2, S3, S4 and S5. The simplified phylogeny indicates the diversification of the major eukaryotic groups, indicated by the different pie charts. Species names are formatted as the first letter of genus name followed by the first letter of the species name, as they appear in supplementary tables (e.g. *Homo sapiens* is abbreviated as Hs). Filled sections indicate presence of the protein, while numbers on the segment indicate how many paralogues of that protein were identified.

**Table 1.** Summary of seipin/sei-1 queries used in similarity searching. The NCBI accession number for the protein sequences are provided, as well as the scientific name of the organism and the associated supergroup/phylum.

| Accession | Organism | Supergroup / Phylum | N-terminal extension | pre-TM domain length | Mechanism of variance |
| --- | --- | --- | --- | --- | --- |
| NP_013508.3 | *Saccharomyces cerevisiae* | Fungi / Ascomycota | No | n/a | n/a |
| NP_850230.1 | *Arabidopsis thaliana* | Archaeplastida / Anthophyta | Yes | 235 | gene duplication |
| NP_174269.1 | *Arabidopsis thaliana* | Archaeplastida / Anthophyta | Yes | 191 | gene duplication |
| NP_197150.1 | *Arabidopsis thaliana* | Archaeplastida / Anthophyta | No | 25 | gene duplication |
| NP_00137295 | *Homo sapiens* | Metazoa / Chordata | Yes | 90 | alternative translation initiation |
| NP_116056.3 | *Homo sapiens* | Metazoa / Chordata | No | 26 | alternative translation initiation |
| NP_00112417 | *Homo sapiens* | Metazoa / Chordata | No | 26 | alternative translation initiation |
| NP_00137295 | *Homo sapiens* | Metazoa / Chordata | Yes | 90 | alternative translation initiation |
| NP_032170.3 | *Mus musculus* | Metazoa / Chordata | Yes | 90 | alternative translation initiation |
| NP_00127775 | *Mus musculus* | Metazoa / Chordata | No | 26 | alternative translation initiation |
| NP_570012.1 | *Mus musculus* | Metazoa / Chordata | No | 26 | alternative translation initiation |
| NP_741547.1 | *Mus musculus* | Metazoa / Chordata | No | 26 | alternative translation initiation |
| NP_570012.1 | *Drosophila melanogaster* | Metazoa / Insecta | No | 57 | n/a |
| NP_741547.1 | *Caenorhabditis elegans* | Metazoa / Nematoda | No | 11 | n/a |

Further, we identified that retention of multiple seipin paralogues or isoforms is also a trait distributed across eukaryotes. Genome annotation has identified multiple seipin isoforms in Mammalia^27,28^ and multiple seipin paralogues in *A. thaliana*^29^, while the single-celled eukaryotes *S. cerevisiae*^26^ and earlier-diverging Metazoa such as *D. melanogaster* and *C. elegans* retain a single copy of seipin^30,31^. This distribution may be consistent with duplication of genes in so-called “higher” eukaryotes with more complex body plans, like the eukaryotic distribution of Hox genes^32^. However, our results indicate that organisms across eukaryotes possess multiple paralogues of seipin, including other single-celled eukaryotes from diverse phyla such as *Stentor coeruleus* (Alveolates), Gephyrocapsa *huxleyi* (Haptophytes), and *Phytopthora infestans* (Stramenopiles) (Figure 2). We also identified multiple paralogues of seipin in other multicellular eukaryotes such as *Chara braunii* (Charophycae), *Trichoplusia ni* (Insecta), *Oryza sativa* (Archaeplastida), and *Patria miniata* (Echinodermata) (Figure 2). Overall, of the 43 genomes in which homologues of seipin were confidently identified, multiple paralogues were identified in 15 (34%) (Figure 2).

### Distribution of N-terminal extensions across eukaryotic diversity

Previous work had noted the presence of two distinct seipin isoform types in Mammalia^2^; alternative translation initiation resulted in those with an “N-terminal extension”, defined as an extended N terminus before the first transmembrane domain (Figure 1), and those without the extension. One study explored whether there is a functional consequence of this N-terminal extension by overexpression of the short isoform in *M. musculus*^33^, but most of the literature in Metazoa treats the various transcripts of seipin arising from alternative splicing and initiation of BSCL2 as functionally equivalent. *Arabidopsis thaliana*, the model plant, despite being extremely evolutionarily distant from the Mammalia, has three paralogues of seipin, one of which also exhibits an N-terminal extension^34^. In *A. thaliana* this trait is retained via gene duplication rather than alternative translation initiation / splicing (Table 1) and there is evidence that it modulates protein-protein interactions between seipin and VAP27, a membrane trafficking protein responsible for ER-cystoskeleton interaction^35^. When the results from our analysis of seipin diversity indicated that multiple seipin paralogues were common and these paralogues often retained N-terminal diversity (Figure 2), we explored potential functional differences between these domains.

We used InterProScan^36^ to annotate the length of the N-terminus of each seipin gene using the first annotated transmembrane domain as a reference point (Table 2, Table S5). We found that there was substantial diversity in the N-terminus length across eukaryotic diversity, even in organisms which retain only a single copy of seipin; sequence length before the first transmembrane domain varied from 4 to 235 amino acids (Table 2, Table S5). Notably, there were multiple organisms, again from supergroups across eukaryotic diversity, that appear to retain multiple copies of seipin with substantially variable N-termini (Table 2). As well as those previously identified in *H. sapiens, A. thaliana*, and *M. musculus*, this variation was also observed in organisms outside the Mammalia from within the same supergroup(s) (*C. braunii, Oryza sativa, Salmo trutta, Spiroglea musicola*), and organisms from other supergroups (*S. coeruleus* (Alveolates), *G. huxleyi* (Haptophytes)). The InterProScan results did not consistently identify a conserved function or domain for the pre-TM N-terminal region, though “signalling peptide” was the most common functional annotation (15% of homologues). 22.5% of homologues also had some annotation in the N-terminal region for intrinsically disordered sequence (Table 2, Table S5).

### Evolutionary trajectory of seipin evolution across eukaryotic diversity

A pan-eukaryotic distribution of a protein suggests homologous descent from a common ancestor. To investigate the evolutionary trajectory of seipin, particularly in lineages with multiple paralogues, across eukaryotes, we used phylogenetic analysis to determine the relatedness of the various homologues of seipin. Our results showed that the relatedness of these genes resembled the patterns of relatedness across eukaryotes (Figure 3), indicative of common descent from an ancestral protein. Gene / transcript duplications appear to be lineage-specific; the only exception is *A. thaliana* and *O. sativa*, where the duplication of the seipin gene into a long and short isoform appears to predate the monocot/dicot split. The long isoform of *A. thaliana* appears to have duplicated again after the split.

**Figure 3.**
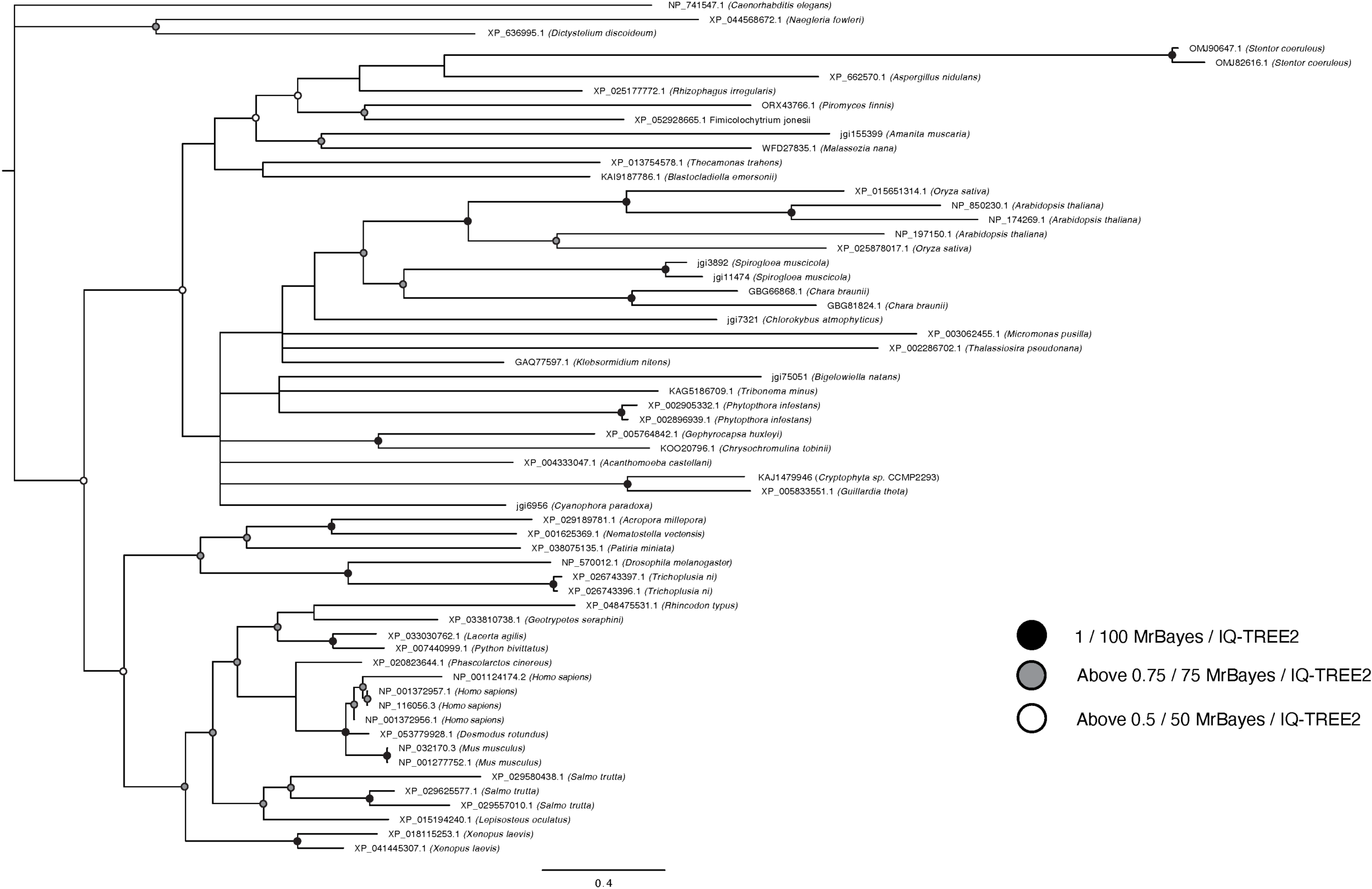
Phylogenetic tree depicting the distribution of seipin across eukaryotic diversity. In this, values for supported nodes (MrBayes/IQ-TREE-2) have been replaced by symbols: black circles 1/100, grey circles >.75/75, and white circles >0.5/50. Node support values are overlaid onto the Bayesian tree topology; where a node is not shared between the Bayesian and Maximum Likelihood support values, this is indicated by a dash where the ML value would be.

### Uncertain evolutionary history of sei-1 in Saccharomyces cerevisiae

A notable result arising from the comparative BLAST results was the lack of observed homology between the previously described functional homologue of seipin in *S. cerevisiae* and any other homologue of seipin outside of the genus *Saccharomyces*. The queries used in the comparative genomic analyses were able to confidently identify seipin homologues in other fungal species (*Aspergillus nidulans, Malassenzia nana, Blastocladiella emersoni, Fimicolochytrium jonesii, Rhizophagus irregularis, Piromyces finnis*), including *A. nidulans*, a species from the same phylum as *Saccharomyces spp.*, but no combination of fungal or non-fungal queries or searches were able to establish a link between the seipin homologues identified in this study and the gene previously described as seipin in *S. cerevisiae*, sei-1. The searches of seipin into *S. cerevisiae* did suggest potential homology to two other *S. cerevisiae* protein families, PFA5 and midasin, albeit with very low subject coverage and a low, though still significant, e value.

To determine whether there was a potential relationship between sei-1 / seipin, PFA5, and midasin, we undertook a phylogenetic analysis of all three proteins in a pan-eukaryotic dataset including all the species with previously annotated copies of seipin (Figure S1). This analysis showed that, despite its divergence from canonical seipin, the sei-1 gene did not group conclusively with either of the other protein families or with seipin. Further phylogenetic analysis of known seipin isoforms compared to a more comprehensive complement of fungal genomes (Table S6, S7) determined that the sei-1 clade did branch within examples of canonical seipin within fungi, but with a tight, highly divergent grouping (Figure 4). These results suggest that sei-1 is a seipin homologue that has undergone accelerated evolution in the Saccharomycetes in particular. However, many other fungal species, including other Ascomycetes such as the model fungus *Aspergillus nidulans*, had canonical seipin paralogues (Figure 4). The divergence in Saccharomycetes appears to be localised to this taxonomic family.

**Figure 4.**
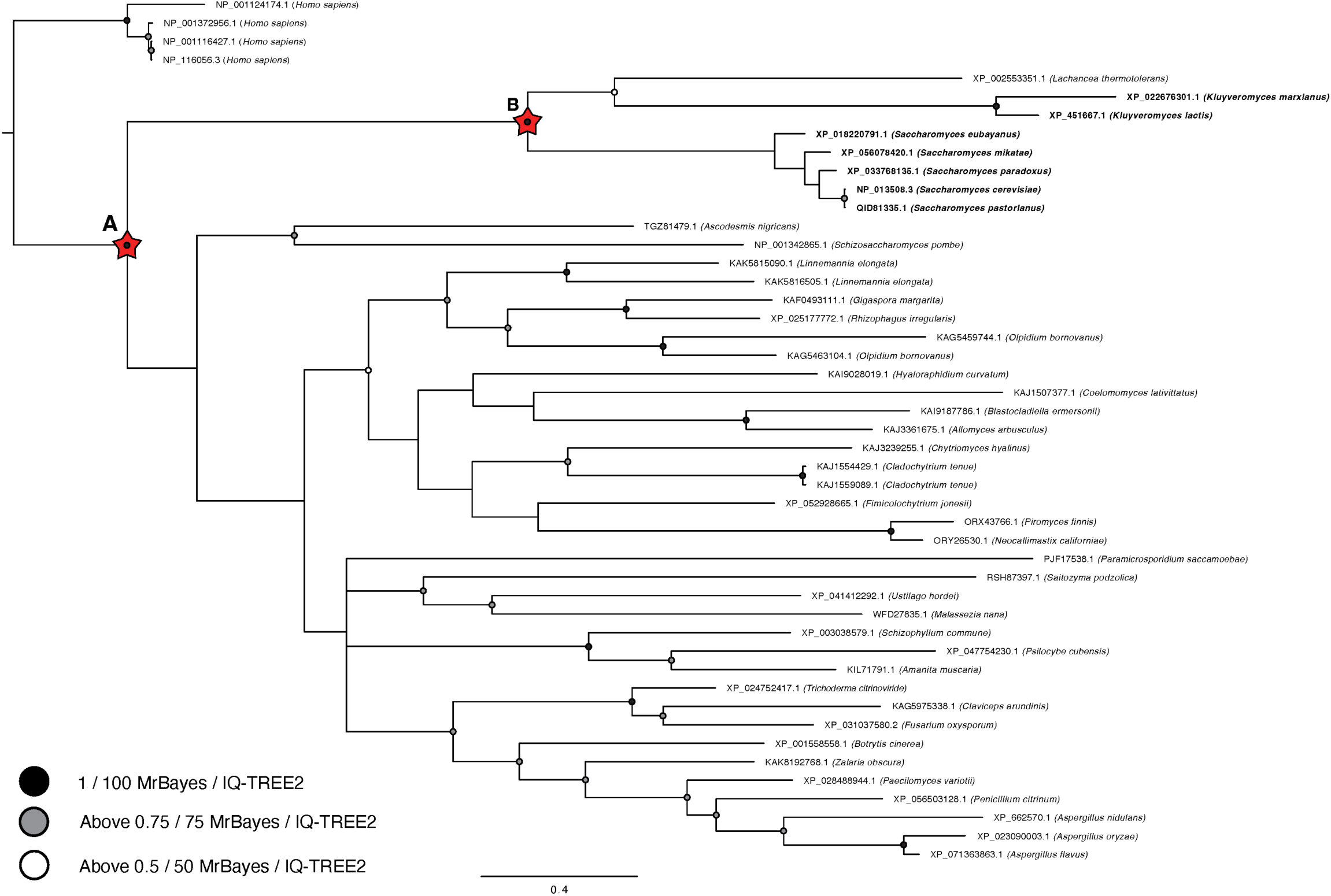
Phylogenetic investigation assessing the distribution of seipin across fungal diversity. *Homo sapiens* seipin isoforms are included as an outgroup. The values for supported nodes (MrBayes/IQ-TREE-2) have been replaced by symbols: black circles 1/100, grey circles >.75/75, and white circles >0.5/50. Red starred nodes indicate evidence for major evolutionary transitions; while sei-1 does group with other fungal seipin sequences (A), they do so with a long-branching node and apparent later sequence divergence within the Saccharomycetes (B). Node support values are overlaid onto the Bayesian tree topology; where a node is not shared between the Bayesian and Maximum Likelihood support values, this is indicated by a dash where the ML value would be.

## Discussion

Previous research had already determined that seipin was likely pan-eukaryotic; it has been comprehensively characterized in model systems in the disparate eukaryotic supergroups Opistokonta and Archaeplastida, which pointed to an ancient origin^1^. This work has substantially increased our understanding of the eukaryotic distribution of seipin across all examined eukaryotic supergroups and lends further support to an evolutionary origin as far back as the Last Eukaryotic Common Ancestor (LECA). Based on functional studies in *Arabidopsis thaliana* and Metazoan model organisms like *Drosophila melanogaster, Mus musculus* and *Homo sapiens*, seipin’s role in LD biogenesis and maintenance is likely conserved across eukaryotes^9,18,26,34^. The extent to which additional functional roles for seipin, and LDs more generally, like those in Metazoa persist in other organisms remains to be determined.

The fact that multiple isoforms and / or paralogues of seipin are maintained across eukaryotic diversity is often indicative of early functional diversification; according to the organelle paralogy hypothesis, duplication of genes is often associated with increased complexity and rapidly emerging dual roles, leading to further specialization of protein families^21^. However, this is not entirely congruent with the phylogenetic evidence, which shows lineage-specific diversification. Lineage-specific diversification, where gene duplication occurs after speciation, is usually more associated with localised functional expansion^37^. The pattern observed with seipin, where gene duplication is both widespread and localised, is unusual and suggests that, over evolutionary time, the role of seipin has diversified multiple times. Some of these duplications are doubtless associated with non-functional genome expansion, such as whole-genome duplications. These have been recorded in numerous organisms, including several of the species used in this study (e.g. *Paramecium tetraurelia*, *Salmo trutta*, and *Xenopus laevis*^38–40^) and we do see expanded seipin complements in some of these organisms. However, we cannot attribute all the observed expansion to non-functional effects. For example, the distribution of seipin in *A. thaliana* and *Oryza sativa* suggests that some of the duplications are ancestral, while others are more recent, outside of known patterns of genome duplications in these species.

Molecular work in *H. sapiens*, *Mus musculus* and *A. thaliana*, where LDAPs are better characterized, suggest that all isoforms / paralogues of the proteins are expressed and have an impact on function^2,25,35^, while an additional copy of a gene arising from whole genome duplication is less likely to be so conserved. It is likely that LECA contained at least one copy of seipin^1^, and its functional role has expanded at multiple places in the evolution of eukaryotes as the role of lipid droplets has also expanded. Further genome sampling to increase the taxonomic resolution of these results in lineages of interest for applied research may be beneficial to fully understand the extent to which LDAPs can be compared between species of interest and model organisms, such as for algal biofuel production. It is also worth noting here that, like with all comparative genomic studies, detection of functional seipin diversification is biased toward organisms which have been studied more intensely, such as *H. sapiens* and *M. musculus*. We examined the protein coding sequences of each species, which in organisms with a rich history of functional work are more likely to include well-annotated isoforms^27,28^. However, organisms for which we had to rely on automated annotations are more likely to under-detect seipin paralogues or isoforms arising from alternative splicing / translation initiation.

It has previously been noted in the literature that seipin maintained distinct isoforms in model organisms that differed primarily in the length of the N-terminal region (defined here as the section of the protein between the N terminus and first transmembrane domain). We sought to identify the distribution and putative function of this N-terminal extension, as we hypothesized that this region may be related to the conservation of multiple seipin paralogues across eukaryotic diversity. We found that, like seipin itself, the variable N-terminal region appears to have arisen multiple times in eukaryotic evolution. This may explain why we found little similarity in length, functional annotation or structure within the N terminus of the genes we analysed (Table 2, S5). However, though there are few functional studies of the N-terminal extension, the existing research does suggest that both proteins are expressed and functional, and knockouts / overexpression studies that affect one isoform or the other show different physiological effects^33,35^. Examining the morphological impact of removing either isoform in *S. coeruleus* may be particularly instructive, as these organisms are from a different supergroup to the previously examined model organisms, exhibit multiple paralogues of seipin with variable N-terminal length, and are amenable to laboratory culturing and genetic manipulation.

Another surprising result from the seipin analysis was the lack of sequence homology between seipin and the *S. cerevisiae* functional homologue of seipin, sei-1. In the pan-eukaryotic BLAST analysis of seipin where sei-1 was used as a query, it failed to retrieve any sei-1 homologues in any other genome and was not retrieved by any of the other query sequences (Table S2, S3). Generally, this level of sequence divergence suggests two proteins are not homologous. However, there is a great deal of evidence from the literature that seipin and sei-1 are at least functionally homologous^26^, and the crystal structures and molecular investigation of both proteins shows a similar domain structure, protein complex structure, and cellular localisation^23,41^.

We therefore sought to determine whether the similarity between seipin and sei-1 was a result of convergent evolution, as initial BLAST queries into the S. cerevisiae genome using known seipin homologues from *H. sapiens, M. musculus, D. melanogaster, C. elegans* and *A. thaliana* detected weak homology to the *S. cerevisiae* genes PFA5 and midasin. However, phylogenetic analysis of sei-1 alongside midasin, PFA5 and seipin genes from representative species failed to determine any conclusive relationships (Fig. S1). To explore this relationship further, we identified seipin / sei-1 homologues from a more taxonomically detailed sampling of fungal clades. This tree, illustrated in Figure 4, was able to demonstrate that the sei-1 genes within the genus Saccharomyces formed a long-branching, monophyletic clade within the detected sei-1 homologues across fungal diversity. While sei-1 does appear to be a highly derived form of seipin, the reason for the rapid diversifying evolution within this lineage specifically remains unknown. It is also notable that many fungal species, including those relatively closely related to *S. cerevisiae*, appear to retain a canonical seipin homologue (Figure 2, Table S2, S3). We did not identify any organisms with both seipin and sei-1, though the yeast species *Lachancea thermotolerans*, also within the family Saccharomycetaceae, has a gene that can be retrieved using seipin queries but groups phylogenetically with the sei-1 genes of other Sacchaomycetes. However, sei-1 queries do not retrieve the *L. thermotolerans* seipin homologue. While this evidence does suggest that seipin and sei-1 are distantly homologous, it may be necessary to exercise caution when inferring seipin function from studies in *S. cerevisiae* / sei-1. This work considerably expands the known evolutionary and functional landscape of seipin, reinforcing its ancient origin and conserved role in lipid droplet biology while also highlighting the complexity of its diversification across eukaryotes. Our analyses suggest that seipin was already present in LECA and subsequent lineage-specific duplications may have repeatedly provided opportunities for functional innovation. The variable N-terminal extensions in model organisms appear to be physiologically distinct, but further molecular characterisation would be required to determine whether this is a conserved trait. Overall, this work shows that seipin evolution across eukaryotic diversity mirrors the complexity of lipid droplets themselves; while there is a broadly conserved function that appears essential to eukaryotic cell biology, diversification across evolutionary lineages have also resulted in idiosyncratic molecular functions which remain to be fully understood.

## Materials and Methods

### Data collection

For the pan-eukaryotic comparative genomic analysis, we used publicly available annotated genome data. We selected genomes to ensure a representative distribution across eukaryotic diversity, though we included additional taxonomic sampling points within the Metazoa as this group contains many model organisms and is of particular interest to a general audience. Database sources and references from which all genomes were obtained are summarized in Table S1. Seipin query sequences were obtained from the relevant genome, based on the number of annotated copies of seipin discussed in the literature^26–28,30,42^, from *Homo sapiens, Mus musculus, Drosophila melanogaster, Caenorhabditis elegans*, and *Saccharomyces cerevisiae*.

### Homology searching

We carried out initial homology searches using the BLAST suite^43^ as implemented in the NCBI web browser, with default parameters. We used a local install of blast+ version 2.17.0 for manual homology searching. We used the BLASTP (protein - protein) algorithm with the query sequences summarised in Table 1 and genomes summarised in Table S1, with default parameters. We then retrieved the fasta sequences from any significant hits and carried out a reciprocal BLASTP into the genome associated with each original query (Table S2). We used the same query and genome dataset to repeat the analyses using the automated comparative genomics software AMOEBAE^44^, using default parameters (Table S3). To reconcile any conflicting results from the two search methods, we examined the alignment scores reported in the AMOEBAE dataset to determine which sequences were likely a result of mis-annotation or pseudogenic sequences.

### Phylogenetics

We generated phylogenetic trees using a combination of maximum likelihood and Bayesian statistical methods. We aligned the input sequences using default parameters in MUSCLE version 5.3 ^45^ and subsequently trimmed them using trimAI version 1.5^46^, with a minimum conservation value of 0.6 for retention of individual sites. For maximum likelihood analysis, we used IQTREE2 version 2.4.0^47^ with 1000 bootstraps, and for Bayesian analysis we used MrBayes version 3.2.7^48^ with mcmcp ngen= 10000000, relburnin=yes, burninfrac=0.25, printfreq=1000, samplefreq=1000, nchains=4.

### Domain prediction

We performed domain prediction using the web interface of InterProScan^36^, and measured the length from the N terminus to the first predicted transmembrane domain. We used the AlphaFold web interface^49^ to confirm the length of the detected N-terminal regions.

## Acknowledgements

EMK, MK, and HT were all supported by Alberta Innovates – Summer Studentship grants (EK 2024, MK and HT 2025). EMK’s work is currently supported by her position as a master’s student with Dr. Joel Dacks. We gratefully acknowledge this support, alongside logistical and administrative support from Mount Royal University and the University of Alberta. We would also like to acknowledge EMK’s Dungeons and Dragons adventuring party for their assistance with time-intensive stages of data curation.

## Author contributions

EMK carried out pan-eukaryotic seipin analysis and phylogenetic analysis of pan-eukaryotic seipin distribution.

HT and MK carried out phylogenetic analysis of the relationship between seipin and sei-1.

FK assisted with database acquisition and curation of publicly available datasets.

ER developed and supervised the research project.

EMK and ER wrote the manuscript.

