## Supplementary figures and images for "Comparative Genomics of the Lipid Droplet-Associated Protein Seipin Across Eukaryotic Diversity Illuminates An Ancient Origin and Conserved Structural Diversity"

### Figure S1

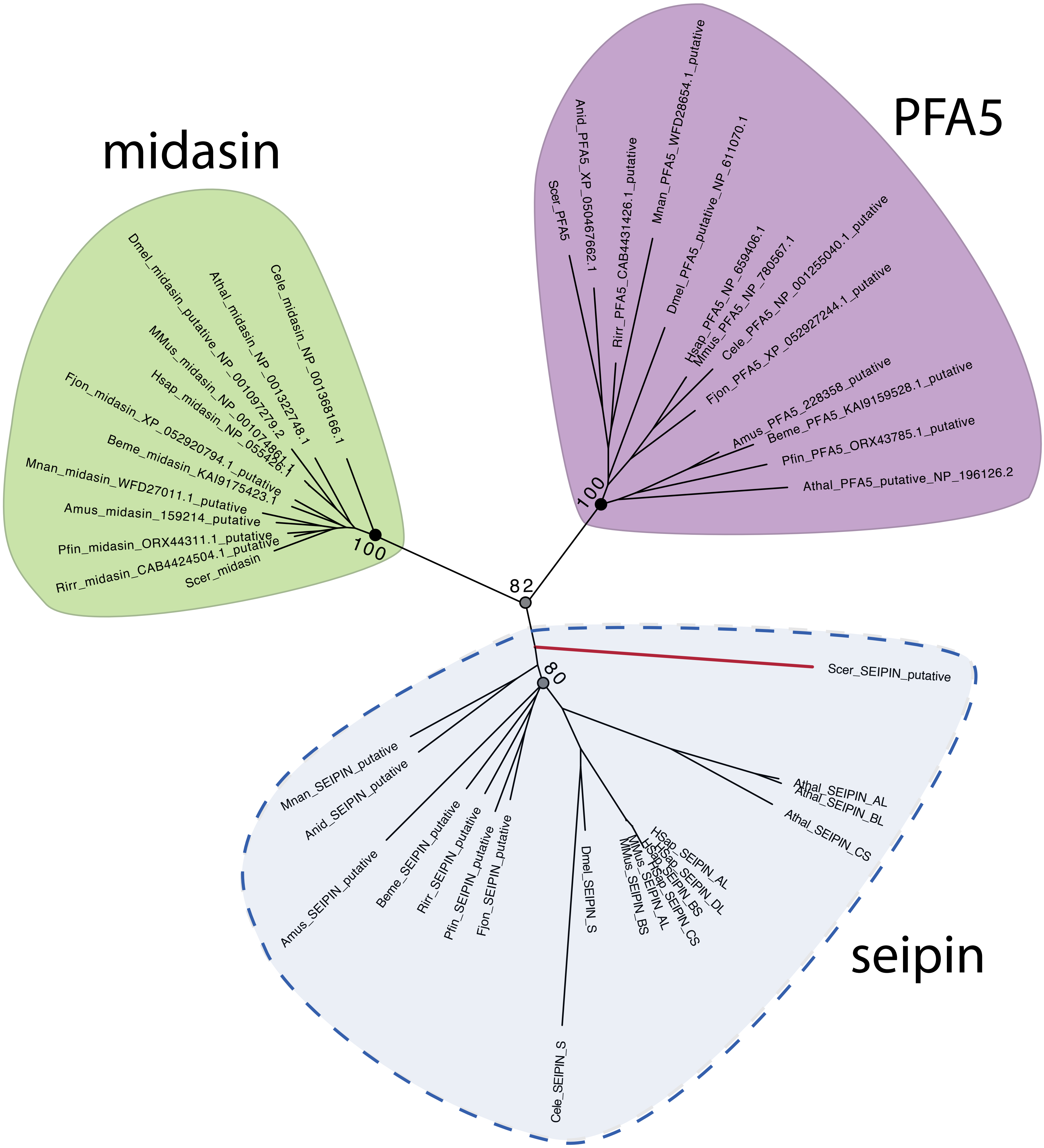
